# Nuclease dead Cas9 is a programmable roadblock for DNA replication

**DOI:** 10.1101/455543

**Authors:** Kelsey Whinn, Gurleen Kaur, Jacob S. Lewis, Grant Schauer, Stefan Müller, Slobodan Jergic, Hamish Maynard, Zhong Yan Gan, Matharishwan Naganbabu, Marcel P. Bruchez, Michael E. O’Donnell, Nicholas E. Dixon, Antoine M. van Oijen, Harshad Ghodke

**Affiliations:** School of Chemistry and Molecular Bioscience and Molecular Horizons, University of Wollongong, Wollongong, New South Wales 2522, Australia; Illawarra Health and Medical Research Institute, Wollongong, New South Wales 2522, Australia; Howard Hughes Medical Institute, Rockefeller University, New York, NY 10065, USA; Department of Chemistry and Center for Nucleic Acids Science and Technology, Carnegie Mellon University, 4400 Fifth Avenue, Pittsburgh, Pennsylvania 15213, USA; Department of Biological Sciences and Molecular Biosensor and Imaging Center, Carnegie Mellon University, 4400 Fifth Avenue, Pittsburgh, Pennsylvania 15213, USA

## Abstract

DNA replication occurs on chromosomal DNA while processes such as DNA repair, recombination and transcription continue. However, we have limited experimental tools to study the consequences of collisions between DNA-bound molecular machines. Here, we repurpose a catalytically inactivated Cas9 (dCas9) construct fused to the photo-stable dL5 protein fluoromodule as a novel, targetable protein-DNA roadblock for studying replication fork arrest at the single-molecule level *in vitro* as well as *in vivo*. We find that the specifically bound dCas9–guideRNA complex arrests viral, bacterial and eukaryotic replication forks *in vitro*.

---

Proteins bound to template DNA can impede DNA replication^1^^-^^3^. Successful replication across such roadblocks requires the coordinated action of several accessory factors and DNA-repair proteins with improper resolution of arrested forks leading to replication fork collapse and, eventually, genetic instability^3^^-^^5^. In addition to lesions and breaks in template DNA, replisomes encounter three major types of protein barriers: transcription complexes, nucleoid-associated proteins, and recombination filaments^6^^-^^8^.

Several tools have been developed to investigate encounters between replication forks and protein barriers. Inspired by the Tus-*Ter* block that terminates replication in *Escherichia coli*, replication fork arrest has been studied at *Ter* sites recombined into the *Saccharomyces cerevisiae* chromosome^9^. Other approaches have involved the introduction of repeat sequences that enable binding of transcription factors to artificially introduce repressor/operator arrays, or proteins that polymerize to form nucleoprotein filaments^10^^-^^13^. Despite their tremendous utility in studying replication fork arrest, these methods suffer from two key disadvantages: (i) tedious recombination techniques are often needed to incorporate tandem arrays of terminator or repressor/operator sequences, and (ii) since the tandem binding of several roadblock proteins is required for effective stalling of the replication fork, the exact positions of the block are often poorly defined. These limitations call for the development of a protein roadblock that is monomeric, binds DNA with high affinity and specificity, and does not require tedious genetic manipulation of template DNA.

We employed the programmable, catalytically inactivated *Streptococcus pyogenes* Cas9 (dCas9) to engineer site- and strand-specific roadblocks to study replication fork arrest at the single-molecule level. To that end, we genetically fused dCas9 to the photostable fluoromodule dL5 that becomes fluorescent upon binding the dye, malachite green (MGE) (Fig. 1a)^14^. First, we purified the dCas9-dL5 fusion protein (Supplementary Table 1, Supplementary Fig. 1a) and assayed its binding to an 83-mer target DNA using surface plasmon resonance (SPR) (Fig. 1b, Supplementary Table 2). Biotinylated target DNA was immobilized on a streptavidin-coated surface and a solution containing dCas9-dL5 pre-programmed with a complementary guide RNA (cgRNA1) was introduced (Fig. 1b, Supplementary Table 3). The dCas9-dL5–cgRNA1 complex exhibited robust and stable binding to the target DNA, whereas dCas9-dL5 alone or in presence of a non-complementary gRNA (ncgRNA) did not exhibit appreciable binding (Fig. 1c, Supplementary Table 2, and Supplementary Note 1). We found that the dCas9-dL5–cgRNA associated strongly and stably with the target DNA – only approximately 25% of the bound complexes dissociated over 16 h (Fig. 1d, Supplementary Note 2).

**Fig. 1:**
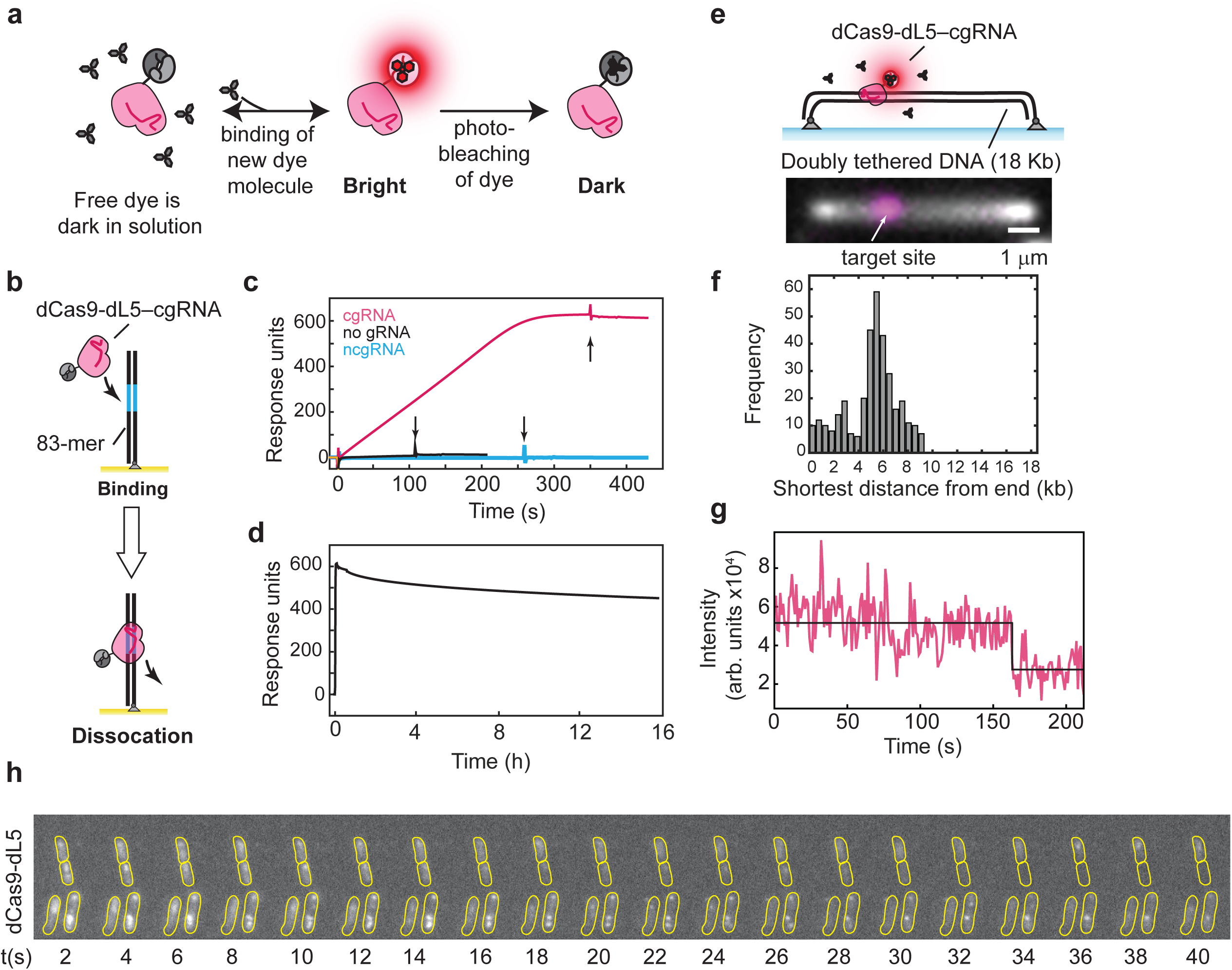
**a.** Schematic of the dCas9-dL5 probe. Binding of MGE enables visualization of dCas9-dL5. P **b.** Schematic of dCas9-dL5 binding to immobilized dsDNA containing the target sequence on an SPR chip. **c.** Sensorgram describing the binding of dCas9-dL5–cgRNA1 to dsDNA substrate carrying the target sequence. Arrows indicate the completion of the injection phase, and switch to running buffer. **d.** Dissociation of dCas9-dL5–cgRNA1 bound to the dsDNA target monitored over 16 h. **e.** Schematic and examples of elongated surface bound and elongated linear dsDNA template bound to dCas9-dL5 (scale bar – 1 μm). dsDNA is stained using Sytox orange, and dCas9-dL5–cgRNA1 is stained by MGE. **f.** Histogram of detected position of dCas9-dL5–cgRNA1 complex visualized by addition of MGE (*n* = 345 molecules). The shortest distance to the position of the dCas9-dL5 is plotted here. **g.** Example photo-bleaching trajectory of dCas9-dL5–cgRNA1–MGE complex (*n* = 345). **h.** dCas9-dL5 targeted to the *recA* locus can be visualized in live *E. coli* cells growing in media supplemented with MGE.

Next, we confirmed that dCas9-dL5 binds specifically to its target sequence. We used single-molecule total internal reflectance fluorescence (TIRF) microscopy to directly visualize dCas9-dL5 bound to its target sequence on individual DNA molecules. DNA molecules were pre-incubated with dCas9-dL5–cgRNA and doubly tethered to a streptavidin-coated glass coverslip inside a microfluidic flow cell using biotinylated oligonucleotide handles (Fig. 1e)^15^. Addition of MGE into the flow cell enabled visualization of the dL5 tag, and positioning of the dCas9-dL5–cgRNA complex along the length of the DNA (Supplementary Note 3; 349 out of 899 templates exhibited bound dCas9-dL5). The position of the bound dCas9-dL5–cgRNA complex was in good agreement with the expected position (Fig. 1f). The use of the MGE allowed us to reliably visualize target-bound dCas9-dL5 for several minutes (Fig. 1g). Taken together, these results demonstrate that the dCas9-dL5–cgRNA1 complex binds with high specificity and stability to its target DNA sequence.

Next, we investigated whether dCas9-dL5 targeted to a specific sequence could be observed in live *E. coli* on long timescales. We expressed dCas9-dL5 from a low-copy number plasmid and targeted it to the template strand of the *recA* gene locus on the chromosome by expressing a cgRNA from a second plasmid (Supplementary Table 1). We then imaged these cells in a flow cell while continuously circulating rich growth medium containing 20 nM of MGE (Supplementary Note 4)^16^. Cells did not exhibit defects in growth rate in the presence of MGE (Supplementary Fig. 1d). Under these conditions, binding of dCas9-dL5 to the target sequence manifested as foci in cells (Figure 1h). The use of the dL5–MGE fluoromodule enabled observation of binding events on a time-scale 15 times longer compared to YPet under conditions of comparable signal to background (Fig. 1h, Supplementary Fig. 1e and 1f, and Supplementary Note 4)^17^. MGE alone did not exhibit foci in cells lacking dL5 (Supplementary Fig. 1g).

We then investigated whether single dCas9-dL5–cgRNA1 molecules bound to template DNA could impede DNA replication using a rolling-circle replication assay, both at the ensemble and single-molecule levels^18^^-^^22^. This assay allows observation of robust DNA synthesis by replisomes under a variety of experimental conditions (Fig. 2a). Pre-incubation of template DNA with dCas9-dL5–cgRNA1 resulted in potent replication fork arrest of reconstituted *E. coli* replisomes during either leading-strand (Fig. 2a and Supplementary Fig. 2a) or simultaneous leading- and lagging-strand DNA synthesis (Supplementary Fig. 2b), with an average blocking efficiency of 85 ± 2% (*N* [replicates] = 5). Importantly, neither complementary gRNA alone (Fig. 2a and Supplementary Fig. 2c) nor dCas9-dL5 alone (Fig. 2a, Supplementary Fig. 2d and Supplementary note 5) or programmed with ncgRNA (Fig. 2a and Supplementary Fig. 2e) could site-specifically arrest DNA replication (summarized in Fig. 2h). Next, to demonstrate the use of this tool in single-molecule assays, we repeated these experiments in single-molecule rolling-circle assays and measured the average lengths of DNA products synthesized by individual *E. coli* replisomes in the presence of dCas9-dL5–cgRNA complexes. Consistent with the bulk experiments, target bound dCas9-dL5 was found to specifically block simultaneous leading- and lagging-strand DNA synthesis (Fig. 2b and Fig. 2c). Further, dCas9-dL5 targeted to the leading strand using a complementary gRNA duplex (cgRNA4 (Ld)) blocked *E. coli* leading-strand and leading- and lagging-strand synthesis with similar efficiencies (85 ± 2% (*N* [replicates] = 5) (Supplementary Fig. 2f). Taken together, these observations demonstrate that the orientation of bound dCas9-dL5–cgRNA on template DNA does not influence its ability to arrest replication.

**Fig. 2:**
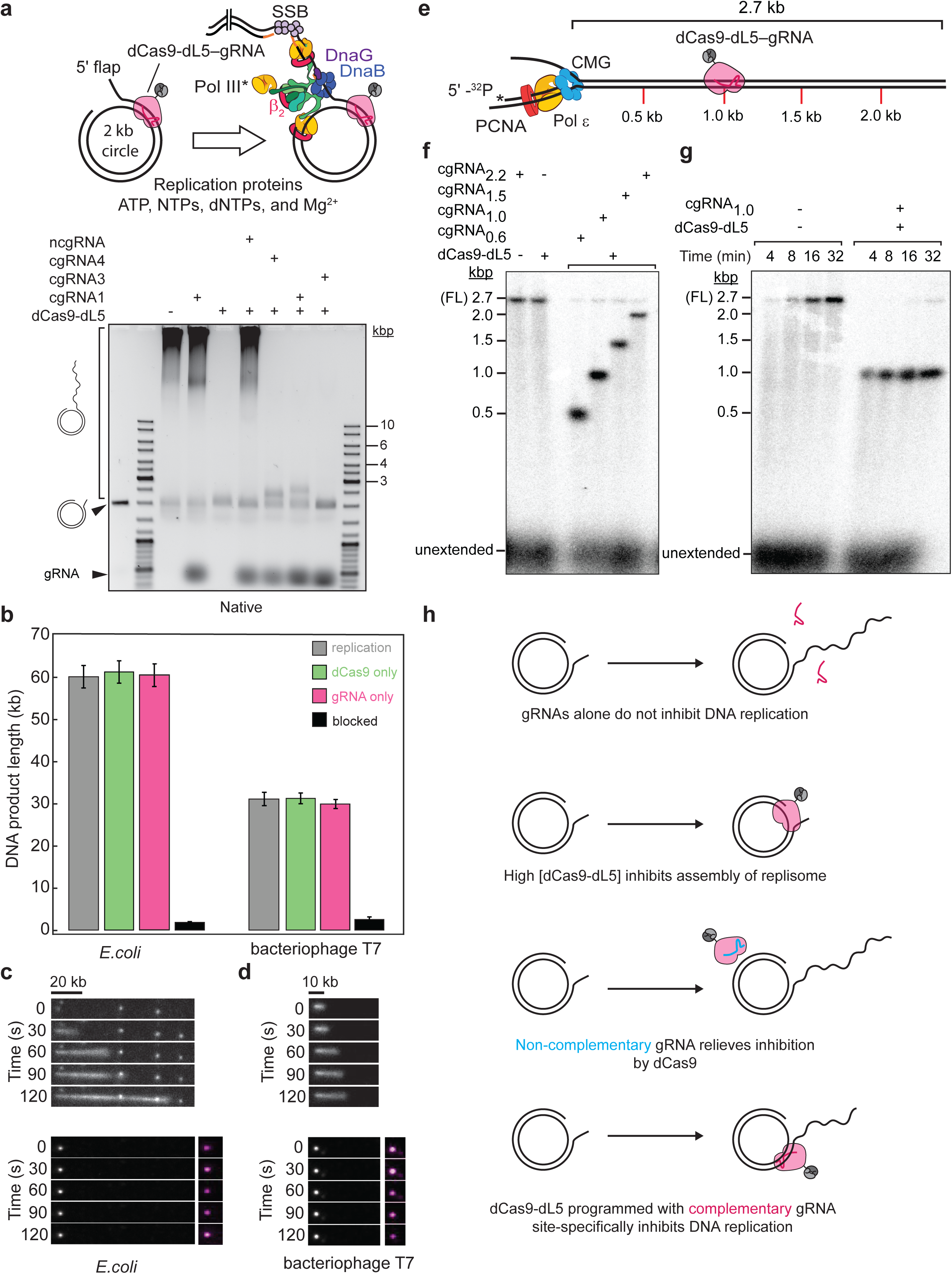
dCas9-dL5 efficiently and stably blocks bacterial, viral, and eukaryotic DNA replication regardless of the targeted strand. **a.** Schematic of the rolling circle DNA replication assay. Addition of the *E. coli* or T7 replication proteins, nucleotides, and Mg^2+^ initiates DNA synthesis. The DNA products are separated by gel electrophoresis by staining with SYBR-Gold, or visualized by single-molecule fluorescence microscopy by staining with Sytox orange. dCas9-dL5 (100 nM) programmed with ncgRNA (400 nM) and cgRNAs alone (400 nM) alone do not inhibit DNA replication. At high concentrations dCas9-dL5 (100 nM) alone inhibits DNA synthesis. See also, Supplementary Fig. 2d for dCas9-dL5 titration. dCas9-dL5 programmed with cgRNAs arrest the progress of the replication fork at the target site. **b.** Bar plots of mean DNA product lengths from *E. coli* and T7 single-molecule rolling circle DNA replication assays. Values plotted are derived from exponential fits to single-molecule DNA product length distributions (*n* > 91 molecules). Error bars indicate errors of the fit. **c.** and **d.** (Top panel) Example kymographs of an individual DNA molecule undergoing DNA replication by *E. coli* (**c**) (*n* = 177 molecules; replication efficiency of 26 ± 2%)and T7 replisomes (**d**) (*n* = 136 molecules; replication efficiency of 24 ± 2%) in the absence of target bound dCas9-dL5. (Bottom panel) Example kymographs of an individual DNA molecule arrested by target bound dCas9-dL5. The gray scale indicates the fluorescence intensity of stained DNA and magenta indicates dCas9-dL5–cgRNA stained by MGE. **e.** Schematic of the eukaryotic DNA replication assay. Eukaryotic replication is blocked by dCas9-dL5 at specific positions on the replication template. **f.** dCas9-dL5 efficiently blocks eukaryotic replication. The sgRNAs used to specifically target the template are indicated. sgRNA_0.6_ and sgRNA_2.2_ block the leading strand; sgRNA_1.0_ and sgRNA_1.5_ block the lagging strand (see supplementary note 6 for exact positions). All reactions were stopped at 16 min. **g.** Time course of eukaryotic replication in the presence or absence of dCas9-dL5 and sgRNA_1000_. Reactions were stopped at 4, 8, 16 and 32 min as indicated. **h.** Summary of interactions of dCas9-dL5 and template DNA. Only the correctly programmed dCas9-dL5-cgRNA complex site specifically inhibits DNA replication.

Finally, we examined the capacity of dCas9-dL5 as a universal tool for arresting replication forks site-specifically; the ability of dCas9-dL5 programmed with complementary gRNA to arrest replication *in vitro* was assessed using model replisomes from T7 bacteriophage (Fig. 2b and 2d) and *S. cerevisiae* (Fig. 2e–g, Supplementary note 6). Replication reactions using both reconstituted replisomes carried out in the presence of template associated dCas9-dL5–cgRNA also exhibited replication fork arrest as observed with *E. coli*.

Here, we have harnessed the specificity and programmability of the CRISPR/Cas9 system and combined it with the photo-stability of the dL5 fluoromodule to repurpose dCas9 as a tool for studying metabolic processes that occur on DNA. Our *in vitro* characterization demonstrates the suitability of the dCas9-dL5 tool for investigating mechanisms that underlie the protein dynamics that govern replication fork rescue at sites of protein roadblocks on template DNA undergoing replication by viral, bacterial, and eukaryotic replisomes. Further, this tool enables investigation of site-specific replication fork arrest in live cells, when programmed with the desired gRNA.

## Author contributions

Conceptualization: H.G.; Methodology: H.G. and J.S.L; Formal Analysis: K.W., G.K., G.S., S.M., J.S.L. and
S.J., Investigation, K.W., G.K., G.S., S.M., J.S.L., H.M., Z.Y.G., H.G. and S.J.; Writing – Review & Editing: H.G., J.S.L. and A.M.V.O.; Funding Acquisition: H.G. and A.M.V.O.; Resources: M.N., M.P.B, N.E.D, M.E. O’D.; Supervision: H.G., J.S.L, A.M.V.O.

## Acknowledgements

HG acknowledges the University of Wollongong and the Faculty of Science, Medicine and Health for SMAH Strategic Equipment funding. A.M.V.O. acknowledges support by the Australian Research Council (DP180100858 and FL140100027). We thank Johan Elf for his gift of the MG1655 dCas9-Ypet strain and Lisanne M. Spenkelink for helpful discussions.

## Supplementary Notes

### Supplementary Note 1: Binding of dCas9-dL5 to dsDNA

We found that highly purified dCas9-dL5 alone exhibited binding to 83-mer biotinylated dsDNA in the absence of guide RNA (Fig. 1c). This minimal binding was lost when dCas9-dL5 was programmed with ncgRNA and may reflect non-specific association of dCas9-dL5 for dsDNA ends.

### Supplementary Note 2: SPR characterization of dCas9-dL5 binding

The sensorgrams obtained during the association of dCas9-dL5-cgRNA at different concentrations of dCas9-dL5 (Supplementary Fig. 1b) exhibited a distinct biphasic profile. The linear response (RU)–time (s) relationship suggests that the association of the complex from solution could be a fast, diffusion limited process. To examine this possibility, solutions of dCas9-dL5 (10 nM) with cgRNA1 (50 nM) in SPR buffer supplemented with 150 mM NaCl and 10 mM MgCl_2_ were injected over immobilized template DNA for 120 s at three different flow rates accessible to the BIAcore T200 instrument: 20, 30 and 80 µL/min (Supplementary Fig. 1c). The observed increase in association rates with the increase in flow rates confirms that the interaction is indeed diffusion-limited, a situation that occurs when the diffusion of analyte from the bulk solution to the chip surface is slower than its binding to the ligand. Conversely, it suggests that the dissociation of analyte (dCas9-dL5–cgRNA) from the ligand (83-mer DNA) must be also be a diffusion limited process, *i.e.* upon dissociation from the ligand, the analyte may not diffuse into bulk solution, allowing it to re-bind. This would result in an apparently slower dissociation rate; therefore, the dissociation half-life (*t*_1/2_) of dCas9-dL5-cgRNA1 from its target dsDNA is an over-estimate of the true dissociation half-life.

### Supplementary Note 3: *In vitro* imaging of dCas9-dL5

Measurement of the position of the bound dCas9-dL5–cgRNA complex on the 18-kb template was performed as follows:

1. Line profiles were manually drawn over all individual DNA molecules. The length of the individual molecules was defined as the distance between the maximum and minimum of the first derivative of the intensity along the drawn lines. Using these measurements, a length distribution was plotted, and values below the 25% and above the 75% percentile were classified as outliers. The resulting distribution was fit to a Gaussian distribution with a mean of 39.5 ± 0.1 pixel. This mean length was then assumed to correspond to the total length of 18,345 bp of the DNA substrate. This conversion resulted in a calibration factor of 466 ± 1 bp/pixel.
2. Next, peaks were detected along the line profile in the MGE-channel. The position of the detected peaks, relative to the ends of the DNA-molecules was then calculated. The distances to both ends of the DNA-molecule were measured. The position in base-pairs was calculated using the calibration described above. The histogram shows the smaller of the two distances from the DNA ends for each molecule.

### Supplementary Note 4: Live-cell imaging of dCas9-dL5

MG1655 cells did not exhibit measurable growth defects when grown in EZ-rich defined medium with glucose as the carbon source supplemented with malachite green dye in the range from 0–200 nM of dye (Supplementary Fig. 1d). Incubation of cells with dye alone did not reveal any non-specific focus formation (Supplementary Fig. 1g). To compare photostability of dL5-MGE to that of YPet, we imaged Mfd-dL5 and Mfd-YPet in cells. Under illumination intensity that yielded comparable signal to background, the time for loss of half the cellular fluorescence signal (Supplementary Fig. 1e) for dL5-MGE was ^~^9 s compared to ^~^0.5 s for YPet.

Supplementary Note 5: dCas9-dL5 alone inhibited rolling circle replication by *E. coli* replisomes at concentrations >20-fold in excess of template DNA concentration (Supplementary Fig. 2d)

In these reactions dCas9-dL5 was pre-incubated with the template DNA that contains a 25-nt single-stranded gap on which the replisome assembles.

### Supplementary Note 6: Ensemble eukaryotic DNA replication assays

Exact positions of targeting gRNA on the eukaryotic linear DNA replication template are as follows: cgRNA_0.6_ targeted nucleotides 583–602 of the leading strand, cgRNA_1.0_ targeted nucleotides 1005–1024 of the lagging strand, cgRNA_1.5_ targeted nucleotides 1493–1512 of the lagging strand, and cgRNA_2.2_ targeted nucleotides 2196–2215 of the leading strand.

**Supplementary table 1.**
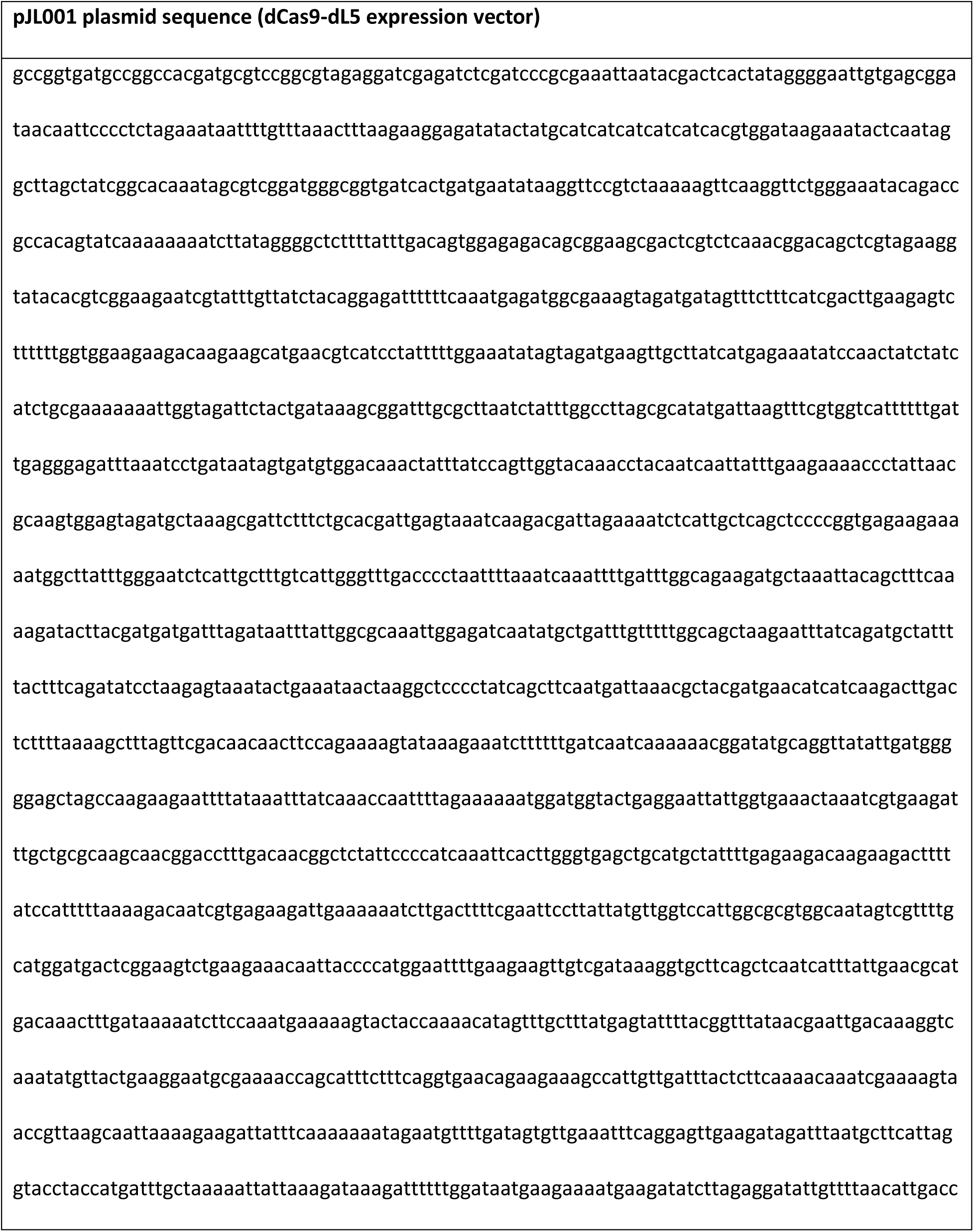
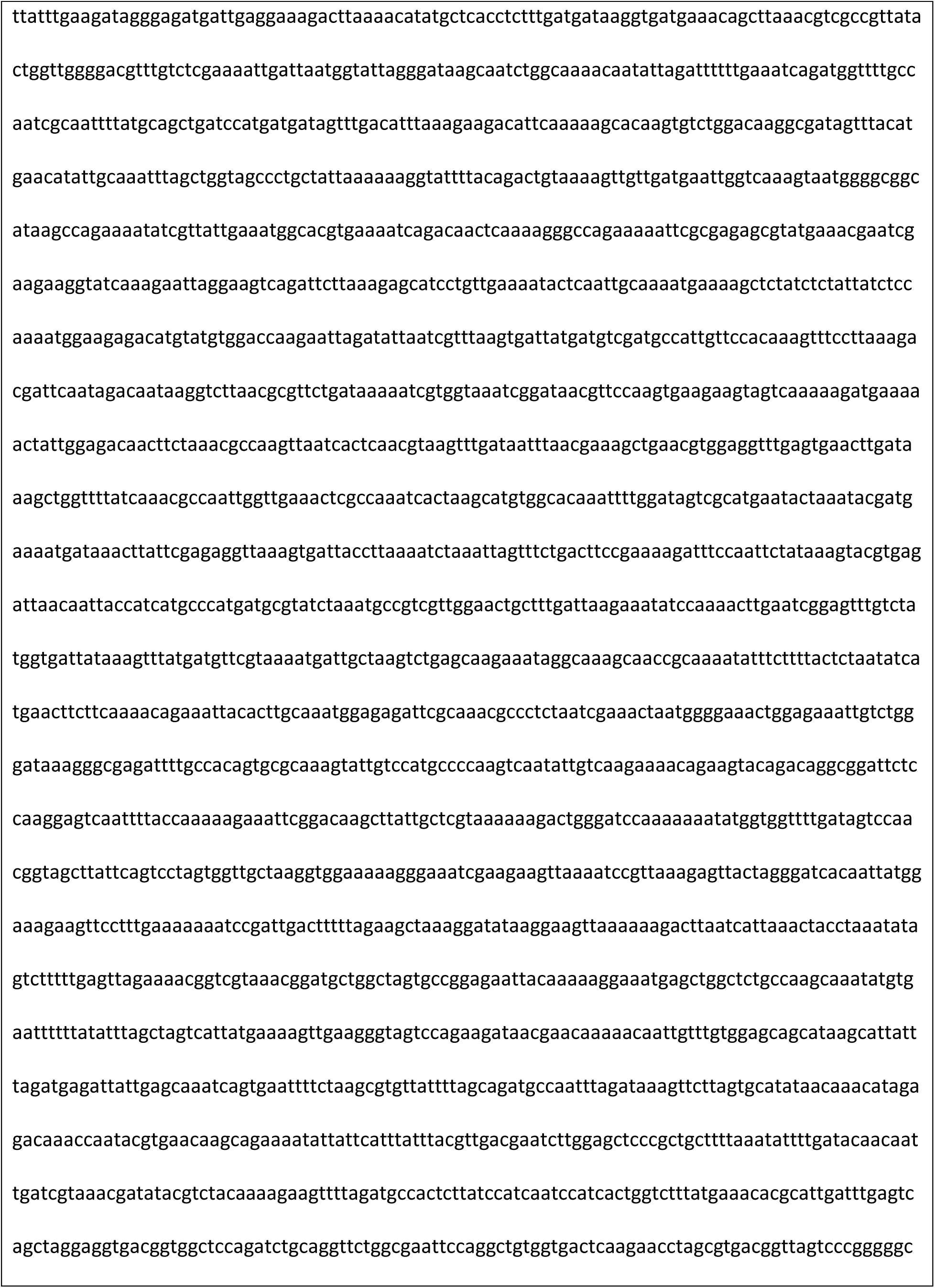
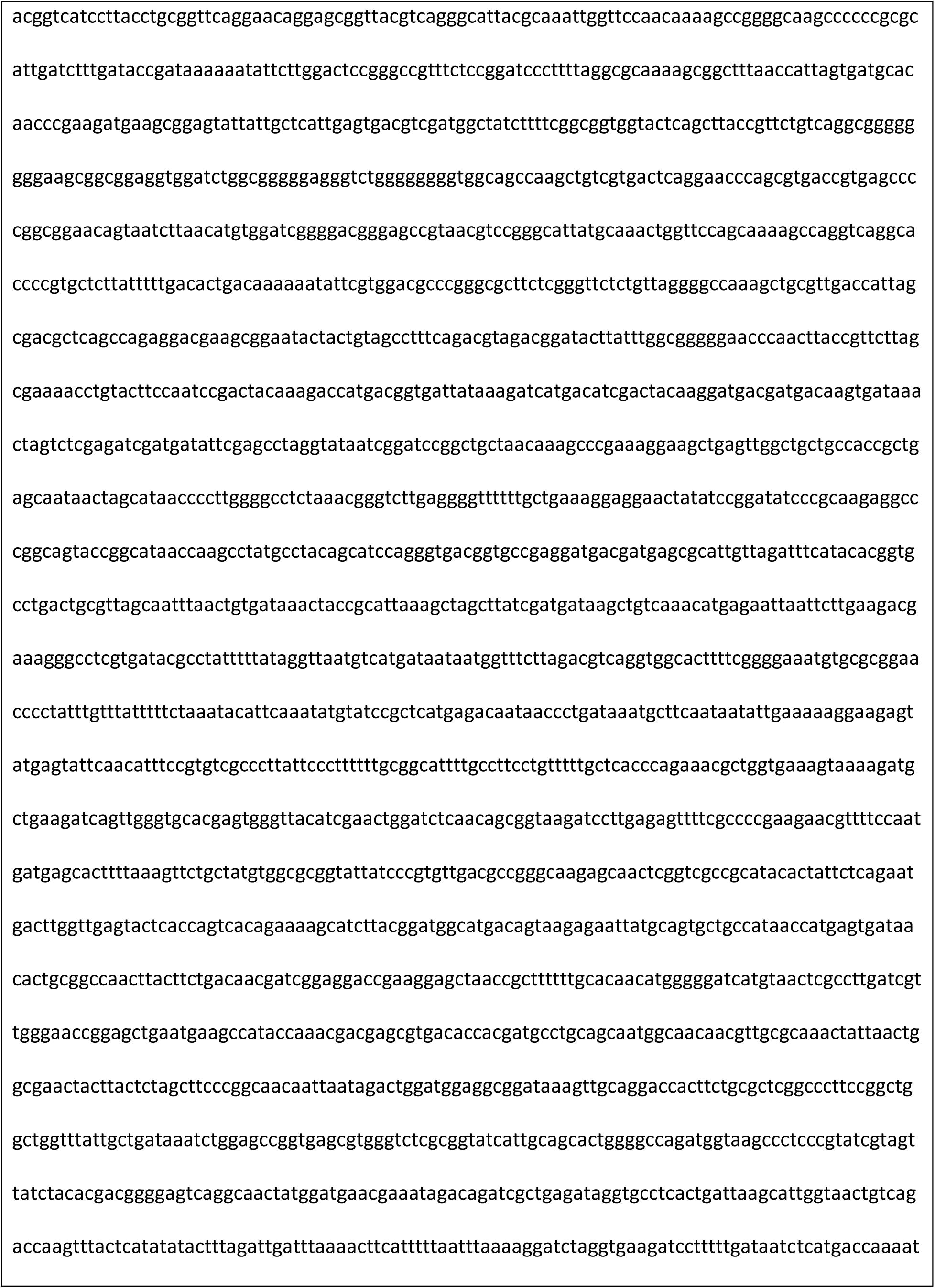
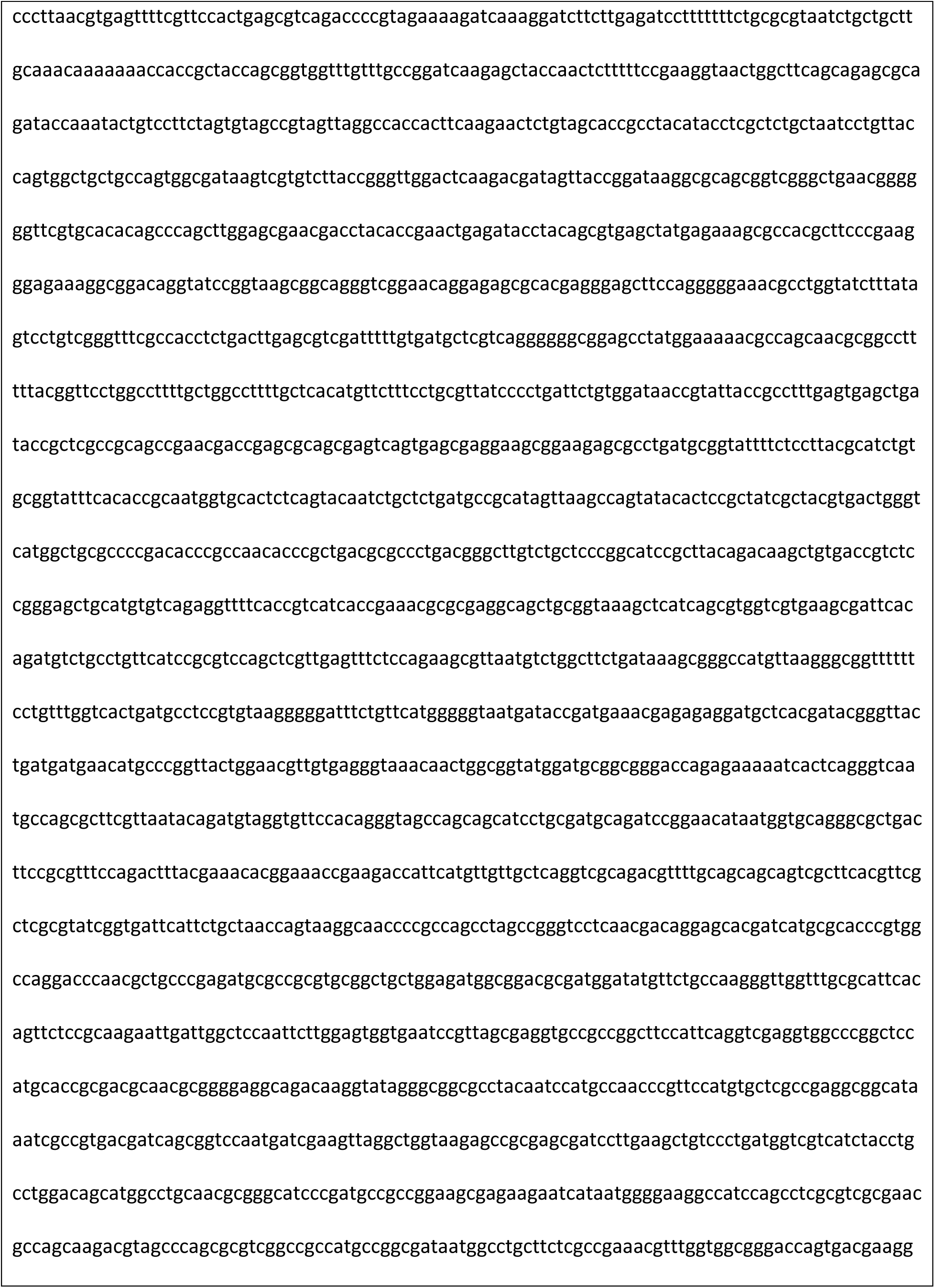
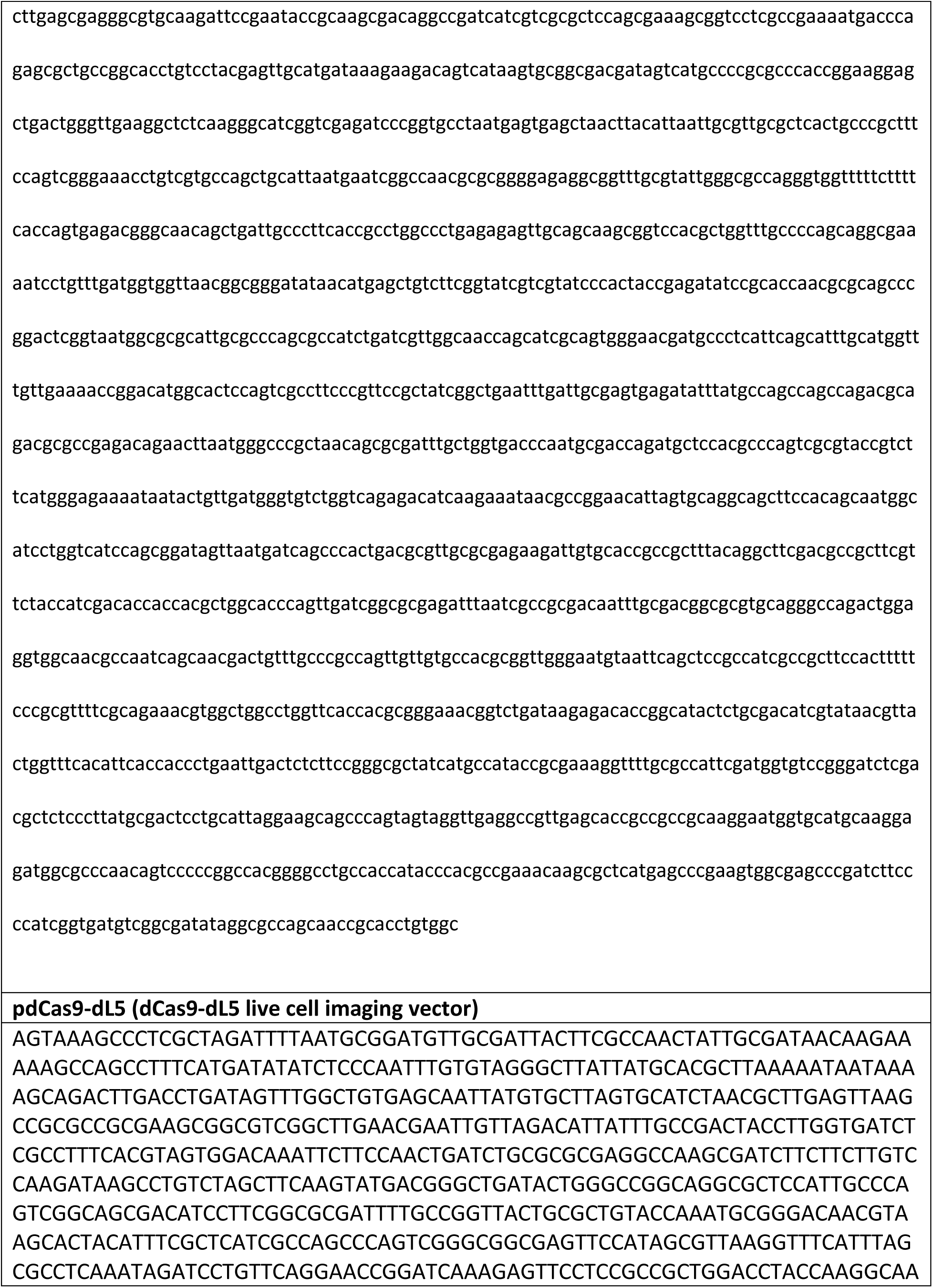
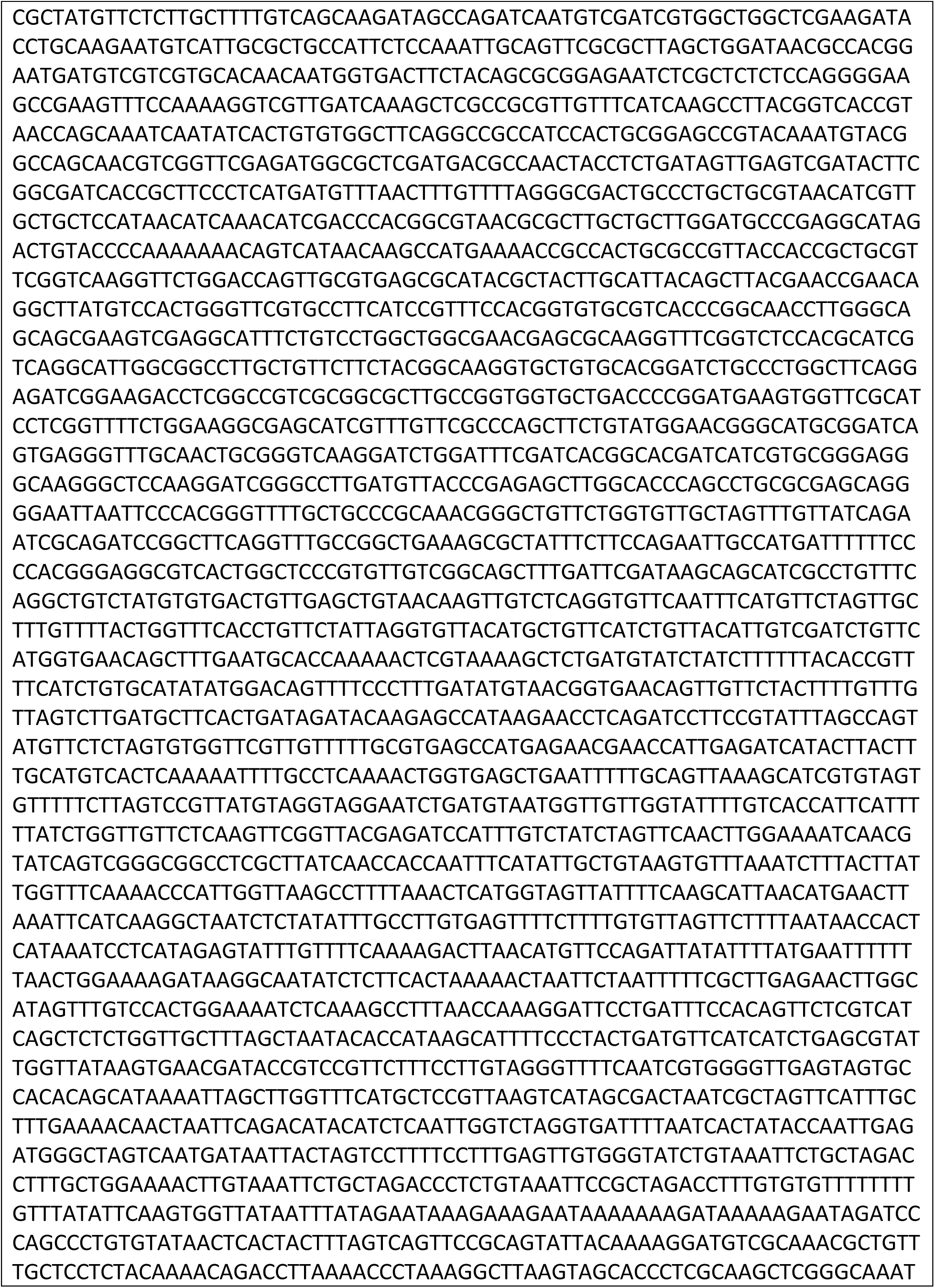
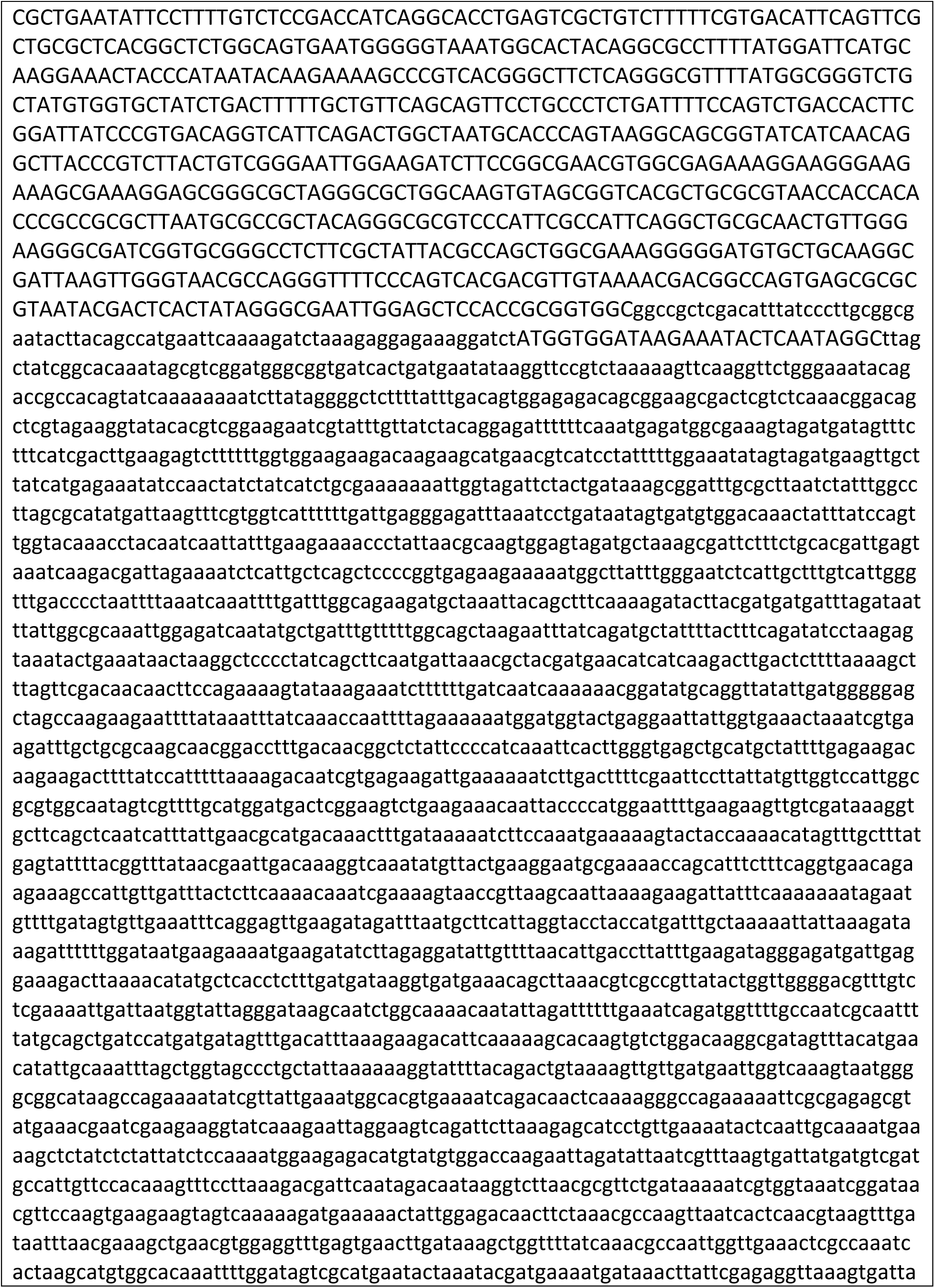
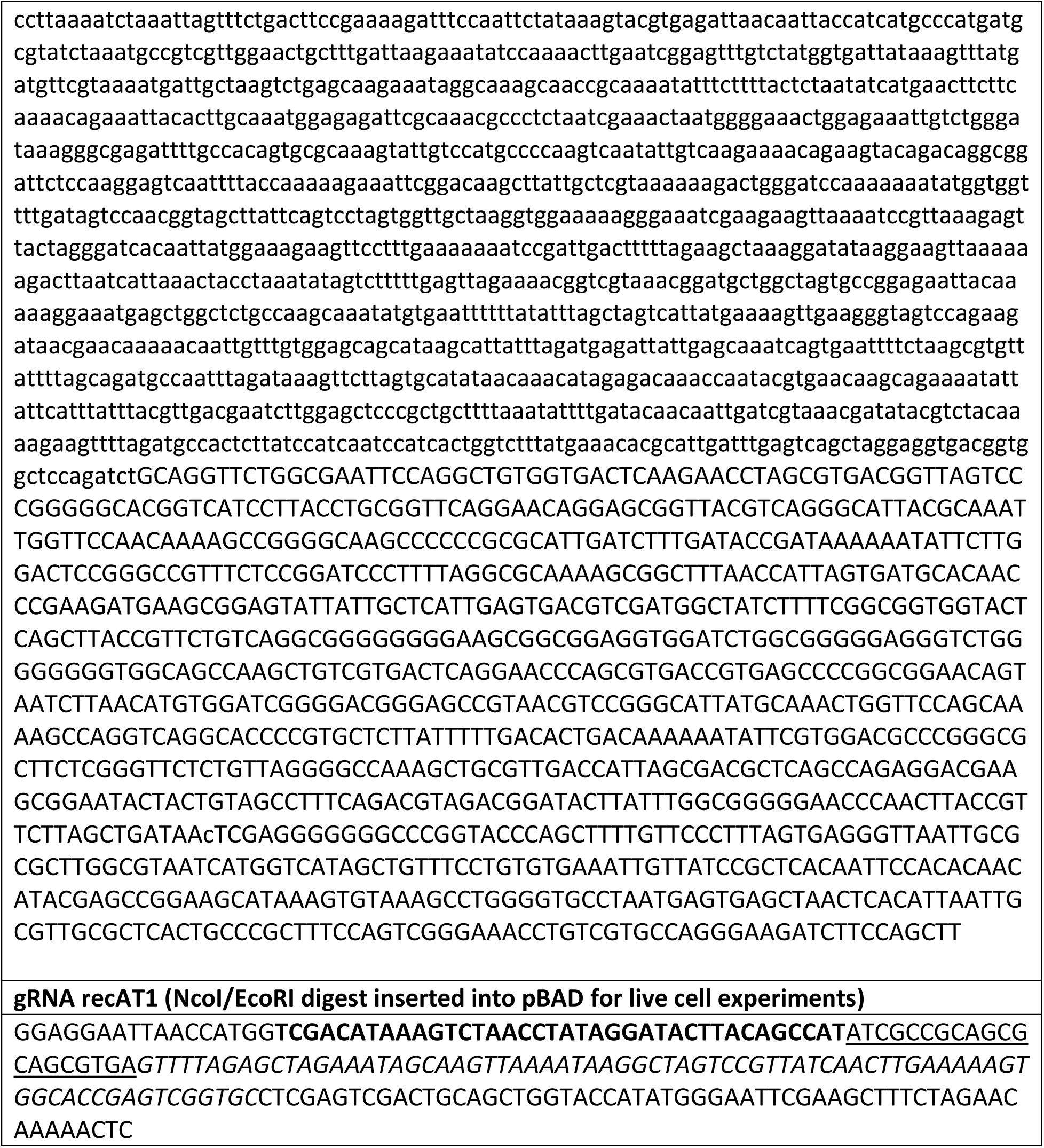

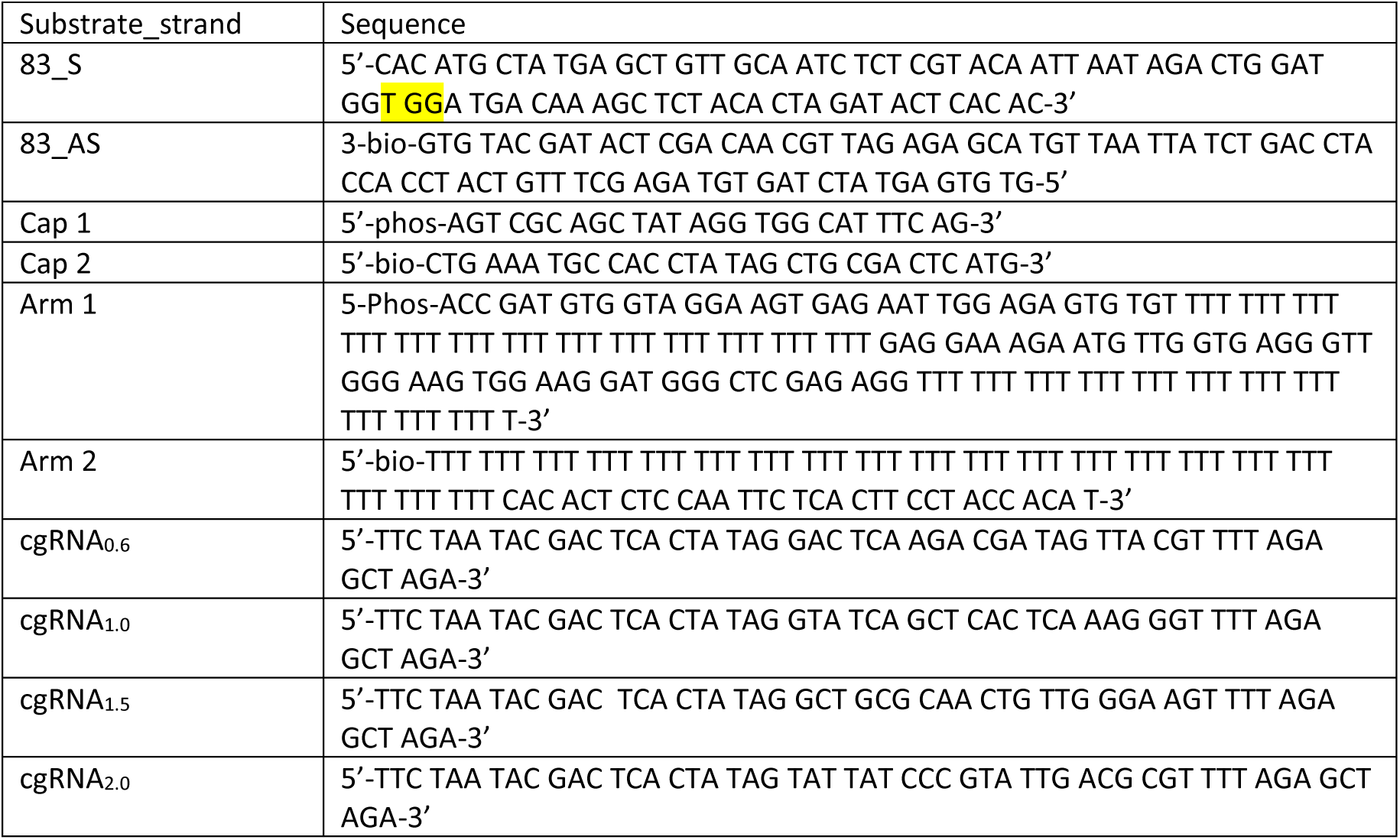

**Supplementary Table 3:**
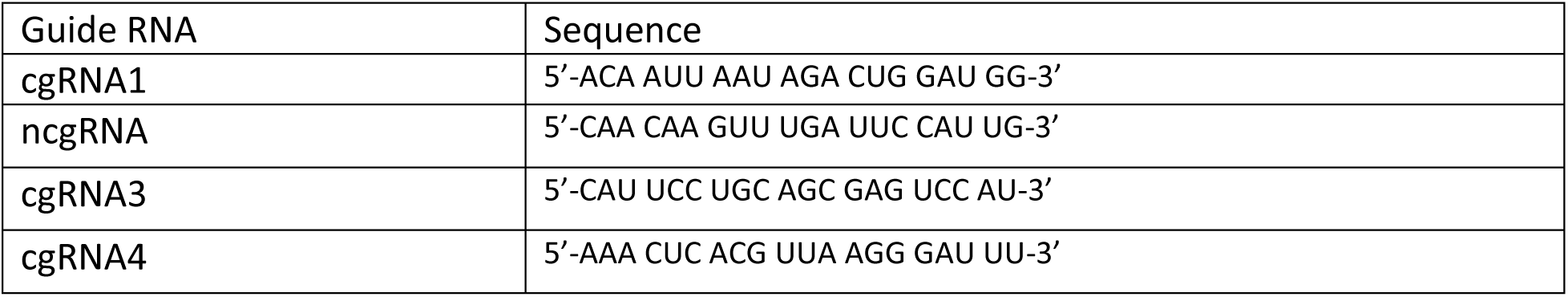
Guide RNA sequences used in this study

## Replication proteins

*E. coli* DNA replication proteins were produced as described previously: the β_2_ sliding clamp^1^, SSB^2^, the DnaB_6_(DnaC)_6_ helicase–loader complex^3^, DnaG primase^4^, the Pol III τ_3_δδ’χψ clamp loader^5^ and Pol III αεθ core^6^. *S. cerevisiae* DNA replication proteins were produced as described previously: the CMG (Cdc45/Mcm2-7/GINS) helicase^7^, the Mrc1–Tof1–Csm3 (MTC) complex^8^, DNA polymerase Pol ε^7^, the PCNA sliding clamp^9^, RPA^7^ and the RFC clamp loader^10^. T7 gp2.5 was produced as described previously^11^. Highly purified T7 gp4 helicase and DNA polymerase gp5/trx were generous gifts of Charles Richardson.

## DNA and RNA oligonucleotides

DNA oligonucleotides and tracrRNA, unmodified crRNAs and crRNAs containing Alexa Fluor 555 were purchased from Integrated DNA Technologies (USA). Sequences of DNA oligonucleotides, crRNAs and tracrRNAs used in this study are listed in Supplementary Table 3. Synthetic guide RNA (gRNA) targeting various regions of the 2.7 kb linear DNA template were produced with the EnGen sgRNA Synthesis Kit (New England Biolabs, USA) using the DNA described in the “DNA used for sgRNA construction” section. All DNA and RNA oligonucleotides were stored in TE buffer (10 mM Tris.HCl pH 8.0, 1 mM EDTA) at –20°C.

## Construction of plasmid pJL001

Plasmid pJL001 was constructed by ligation of a 1007 bp *Sac*I –*Xho*I gene block (Aldervon, USA) between the corresponding sites in pET302 (obtained from Addgene plasmid #72269), to encode dCas9-dL5 containing a N-terminal 6xHis and C-terminal 3xFLAG tags. The corresponding plasmid sequence is given in Supplementary Table 1.

## Expression and purification of dCas9-dL5

*E. coli* strain Rosetta 2(DE3) containing plasmid pJL001 was grown in LB medium supplemented with thymine (25 mg/mL) and ampicillin (100 μg/mL) at 37°C. Upon growth to *A*_600_ = 0.8, the temperature was reduced to 16°C and protein expression induced by addition of 0.5 mM isopropyl-β-D-thiogalactoside. Cultures were further shaken for 16 h at 16°C, then chilled on ice. Cells (8 g from 2 L of culture) were harvested by centrifugation, frozen in liquid nitrogen and stored at –80°C. All subsequent steps were carried out in a cold room maintained at 6°C. After thawing, cells were resuspended in lysis buffer (20 mM Tris.HCl pH 7.6, 0.1 mM EDTA, 1 mM dithiothreitol, 150 mM NaCl, 5% (*v/v*) glycerol) and 2x Protease Inhibitor Cocktail tablets and 0.7 mM phenylmethylsulfonyl fluoride were added to inhibit proteolysis. Cells were lysed by being passed twice through a French press (12,000 psi), and cell debris were then removed by centrifugation. Crude supernatant (85 mL) was brought to 0.4% (*v/v*) in polyethylenimine (PEI) and vigorously stirred. After 40 min, the white precipitate was separated by centrifugation. The remaining pellet was homogenized by stirring in lysis buffer for 15 min. The remaining white precipitate was immediately collected by centrifugation and the supernatant discarded. The remaining pellet was further homogenized in lysis buffer + 250 mM NaCl for 15 min. After centrifugation, the high salt supernatant containing dCas-dL5 was collected yielding Fraction I (72 mL). Proteins that were precipitated from Fraction I by addition of solid ammonium sulfate (0.32 g/mL) and stirring for 60 min, were collected by centrifugation and dissolved in 30 mL of FLAG buffer (25 mM Tris.HCl pH 7.6, 1 mM EDTA, 1 mM dithiothreitol, 200 mM NaCl and 5% (*v/v*) glycerol). The solution was dialysed against 2 L of the same buffer overnight, to yield Fraction II. Fraction II was added to 4 mL FLAG M2 resin prepared as per manufacturer’s instructions and left to incubate with constant mixing. After 1 h, the FLAG M2 resin was poured into a PD-10 column and equilibrated in FLAG wash buffer (50 mM Tris.HCl pH 7.6, 0.5 mM dithiothreitol, 0.5 mM EDTA, 300 mM NaCl, 5% (*v/v*) glycerol). The column was washed with FLAG buffer until the *A*_280_ was approximately 0.05, and dCas9-dL5 was eluted using FLAG wash buffer containing 3X FLAG peptide (200 μg/mL). Fractions containing dCas9-dL5 were collected and pooled to yield Fraction III (15 mL), which was dialysed against 2 L of HisTrap buffer (50 mM Tris.HCl pH 7.6, 0.5 mM EDTA, 2 mM dithiothreitol, 300 mM NaCl, 20 mM imidazole pH 8.0, 5% (*v/v*) glycerol). Fraction III was applied at 1 mL/min onto a 5 mL HisTrap column equilibrated in HisTrap buffer. The column was washed until *A*_280_ returned to baseline and dCas9-dL5 was eluted as a single peak with a step elution of 300 mM imidazole pH 8.0. Fractions under the peak were pooled and dialysed against 2 L of storage buffer (50 mM Tris.HCl pH 7.6,1 mM EDTA, 3 mM dithiothreitol, 300 mM NaCl, 50% (*v/v*) glycerol) to give Fraction IV (4 mL, containing 6.9 mg of protein; Fig. S1a). Aliquots were stored at –20°C.

## Rolling-circle replication template

DNA rolling circle substrates were prepared as previously described^12^.

## Linear DNA substrates

Plasmid pSupercos1 DNA^13^ (7 pmol) was linearized overnight at 37°C with 100 U of *Bst*XI in 1 x buffer 3.1 (New England Biolabs). The 18,345 bp fragment was purified with a Wizard SV gel and PCR clean up kit (Promega) and the concentration was measured. DNA oligonucleotides (750 pmol of arm 1, 4500 pmol arm 2, and 70 pmol capping 1, 2) were annealed by heating at 94°C for 5 min before slow cooling. The biotinylated handles were ligated to the 18,345 bp fragment in 1 x T4 ligase buffer and 2000 U of T4 ligase overnight at 16°C. Biotinylated linear DNA substrates were purified from excess DNA oligonucleotides by adjusting NaCl to 300 mM and loaded by gravity onto a Sepharose 4B (1 x 25 cm) column, equilibrated in gel filtration buffer (10 mM Tris.HCl pH 8.0, 1 mM EDTA, and 300 mM NaCl). Biotinylated linear DNA substrates eluted as a single peak in the column void volume, fractions under the peak were analyzed by agarose gel electrophoresis. Fractions containing linear DNA substrates were pooled and dialysed overnight in 2 L of sterilized TE buffer, concentrated 2-fold in a vacuum concentrator and the concentration measured. This protocol typically yielded ~20 μg DNA. Aliquots were stored at –80°C.

## Forked linear DNA substrates

The DNA replication template, a linearized 2.7 kb plasmid ligated to a synthetic replication fork, was prepared as previously described^7, 14^. A synthetic 37-mer oligonucleotide (Fork primer) was end-labeled with ^32^P-ATP by T4 polynucleotide kinase (New England Biolabs, USA) according to manufacturer’s instructions and annealed to the forked substrate by heating to 85°C and slowly cooling.

## Assessment of dCas9 interactions by surface plasmon resonance (SPR)

SPR measurements were carried out on a BIAcore T200 instrument (GE Healthcare) at 20°C in SPR buffer (30 mM Tris-HCl pH 7.6, 0.5 mM dithiothreitol, 0.25 mM EDTA, 0.005% (*v/v*) surfactant P20) containing NaCl/MgCl_2_ concentrations as described. A streptavidin-coated (SA) sensor chip was activated with three sequential injections of 1 M NaCl, 50 mM NaOH (40 s each at 5 μL/min). Then, a solution (2.5 nM) of the 3’-biotinylated 83-mer template dsDNA in SPR buffer containing 50 mM NaCl (SPR running buffer), assembled *in situ* by premixing 83-S and 83-AS oligonucleotides (to final concentrations of 1.2 and 1 µM, respectively) in hybridization buffer (20 mM Tris-HCl pH 7.8, 50 mM NaCl, 5 mM MgCl_2_) at 90°C for 5 min followed by slow cooling overnight to the room temperature, was used to immobilize ~150 RU of DNA template at 5 μL/min over 456 s onto the surface of flow cell 4, whereas flow cell 3 was left unmodified and served as a control (4–3 subtraction).

To interrogate binding specificity of dCas9-dL5 for immobilized 83 dsDNA template in the presence of complementary guide cgRNA1, a solution of protein (10 nM) with cgRNA1 (50 nM) in SPR buffer supplemented with 100 mM NaCl and 10 mM MgCl_2_ (SPR binding buffer) was made to flow at 10 μL/min for 350 s, yielding a response of ~625 RU (Fig. 1c). Following the association phase, the slow dissociation of protein from immobilized DNA template initiated by re-introduction of the running buffer in the flow cell and monitored over >70 s indicated stable binding. Bound proteins/RNA complexes were removed and immobilized dsDNA on the chip surface regenerated by three successive 40 s injections of 3 M MgCl_2_ at 10 μL/min. Injections of dCas9-dL5 under similar experimental conditions, either in the presence of ncgRNA (257 s injection) or in the absence of any guide gRNA (107 s), as well as the injection of cgRNA1 alone (66 s) led to barely detectable binding responses, suggesting that only the dCas9-dL5–cgRNA1 complex interacts stably and specifically with 83 template dsDNA. Furthermore, binding of dCas9-dL5–cgRNA1 is concentration dependent, since comparative injection of 30 nM dCas9-dL5 with 50 nM cgRNA led to faster association (Fig. S1b). Moreover, notably similar responses measured at equilibrium when 10 nM and 30 nM dCas9-dL5 were injected with 50 nM cgRNA1 (~625 RU) implies saturation of all the template DNA molecules on the chip surface with 10 nM dCas9-dL5–cgRNA1, indicating: (a) that the *K*_D_ for the dCas9-dL5–cgRNA1–dsDNA interaction is significantly below 10 nM in buffer containing 100 mM NaCl and 10 mM MgCl_2_, and (b) that the dCas9-dL5–cgRNA1 complex binds 83-mer template DNA in 1:1 molar ratio, *i.e.* considering that the ratio of mol. wt. between dCas9-dL5–cgRNA1 complex (~218.1 kDa; 184.5 kDa for dCas9 and ~33.6 kDa for cgRNA1) and template dsDNA (51.7 kDa; 25.5 kDa for 83-S and 26.2 kDa for 83-AS) is 4.2, and considering that ~150 RU of DNA was immobilized on the surface, ~630 RU (4.2·150 RU) of bound dCas9-dL5–cgRNA1 could be expected at saturation in case of 1:1 interaction with template DNA.

To demonstrate the strong association and long-term stability of dCas9-dL5–cgRNA1 complex with the target DNA template, the dissociation of a complex assembled on the surface during injection of 30 nM dCas9-dL5 and 50 nM cgRNA1 (as described above) from immobilized DNA in SPR running buffer, interspersed with an early 1500 s injection of SPR binding buffer to assess the complex stability in the buffer used for the association, was monitored for over 16 h (58807 s; final response was ~450 RU; Fig. 1d). The surface (immobilized template dsDNA) was then regenerated with one 40 s injection of 3 M MgCl_2_ at 10 μL/min. Assuming first-order dissociation and SPR responses that were measured following the injection of SPR binding buffer, at the start of measured dissociation *R*_0_ = 575 RU and at the end *R*_t_ = 450 RU over the period of t = 57000 s, the dissociation half-life of > 44 hours (see also Note 2) was calculated using Equation 1 : 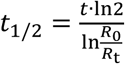

## Measurement of dCas9-dL5 binding specificity on long DNA substrates

Microfluidic flow cells were prepared as described in “Preparation of flow-cells for *in vitro* imaging”. To help prevent non-specific interactions of proteins and DNA with all surfaces, they were blocked with 2% Tween20 in blocking buffer (50 mM Tris-HCl pH 7.6, 50 mM KCl). Imaging parameters are described in “*In vitro* single-molecule fluorescence microscopy”.

First, 9 nM dCas9-dL5 was incubated with 15 nM cgRNA1 at 37°C for 5 min in reaction buffer (25 mM Tris-HCl pH 7.6, 10 mM MgCl_2_, 150 mM potassium glutamate, 0.1mM EDTA and 0.0025% (*v/v*) Tween20). The dCas9-dL5–cgRNA1 complex was further incubated with 125 pM biotinylated linear DNA substrates at 37°C for 20 min in reaction buffer supplemented with 0.5 mg/mL heparin. To reduce heterogeneity in DNA lengths upon binding to the surface, 200 µM chloroquine was added immediately prior to injection of the sample into the flow cell. The solution was injected at a constant rate of 17 µL/min until an appropriate DNA density was achieved. Next, the flow cell was washed with 2 mL of reaction buffer, supplemented with 100 mM NaCl, 15 nM gRNA and 0.5 mg/mL heparin. dCas9-dL5–cgRNA1–DNA complexes were imaged in reaction buffer containing 150 nM Sytox orange and 150 nM MGE.

## Ensemble *E. coli* replication assays

Standard leading-strand replication assays were set up in replication buffer (RB; 60 mM Tris-HCl pH 7.6, 24 mM Mg(OAc)_2_, 100 mM potassium glutamate, 1 mM EDTA and 0.005% (*v/v*) Tween20) and contained 2 nM rolling-circle replication template, specified concentrations of dCas9-dL5 and gRNA, 60 nM DnaBC, 30 nM τ_3_δδ’χψ, 90 nM Pol III αεθ core, 200 nM β_2_, 10 mM dithiothreitol, 1 mM ATP, and 125 µM dNTPs in a final volume of 12 µL. First, dCas9-dL5 was incubated with gRNA for 5 min, and further incubated with rolling-circle DNA templates for 5 min at room temperature. Components (except dCas9-dL5–gRNA–DNA) were mixed and treated at room temperature, then cooled in ice for 5 min prior to addition of dCas9-dL5–gRNA–DNA complexes. Reactions were initiated at 30°C and quenched at specified time points by the addition of 200 mM EDTA and 2% (*w/v*) SDS. The quenched reactions were loaded into a 0.6% (*w*/*v*) agarose gel in 2x TAE. Products were separated by agarose gel electrophoresis, at 60 V for 150 min and stained in SYBR-Gold (Invitrogen, Waltham, MA) and imaged under UV light.

*E. coli* leading- and lagging-strand DNA replication reactions were carried out as previously described^6^ with the following minor modifications. Reactions were set up in RB, and contained 4 nM rolling-circle replication template, specified concentrations of dCas9-dL5 and gRNA, 60 nM DnaBC, 80 nM DnaG, 30 nM τ_3_δδ’χψ, 10 nM SSB, 90 nM Pol III αεθ core, 200 nM β_2_, 10 mM dithiothreitol, 1 mM ATP, 125 µM dNTPs, and 250 µM NTPs to a final volume of 12 µL, quenched after 30 min by addition of 1.5 μL 0.5 M EDTA and 3 μL DNA loading dye (6 mM EDTA, 300 mM NaOH, 0.25% (*v*/*v*) bromocresol green, 0.25% (*v*/*v*) xylene cyanol FF, 30% (*v*/*v*) glycerol). DNA products were separated on a 0.6% (*w*/*v*) alkaline agarose gel at 14 V for 14 h. The gel was then neutralized in TAE buffer, stained with SYBR-Gold and imaged under UV light.

## Ensemble T7 replication assays

T7 leading-strand DNA replication assays were carried out using previously described conditions^15^. Briefly, reactions were set up in T7 replication (TR) buffer (25 mM Tris-HCl pH 7.6, 10 mM MgCl_2_, 50 mM potassium glutamate, 0.1 mM EDTA and 0.0025% (*v/v*) Tween20) and contained 2 nM rolling-circle replication template, specified concentrations of dCas9-dL5 and cgRNA1, 180 nM gp2.5, 5 nM gp4, 40 nM gp5, 10 mM dithiothreitol, 1 mM ATP, 1 mM CTP, and 600 µM dNTPs, in a final volume of 12 µL. First, dCas9-dL5 was incubated with cgRNA1 for 5 min, and further incubated with rolling-circle DNA templates for a further 5 min at room temperature. Components (except dCas9-dL5– cgRNA1–DNA) were mixed and treated at room temperature, then cooled in ice for 5 min prior to addition of dCas9-dL5–cgRNA1–DNA complexes. Reactions were initiated at 30°C and quenched at specified time points by the addition of 200 mM EDTA and 2% (*w/v*) SDS. The quenched reactions were loaded onto the 0.6% (*w/v*) agarose gel, which was run under the same conditions as standard *E. coli* leading-strand replication assays.

## Ensemble *S. cerevisiae* replication assays

Leading-strand replication assays were set up in eukaryotic replication (ER) buffer (25 mM Tris-OAc pH 7.5, 5% glycerol, 80 μg/mL BSA, 5 mM tris(2-carboxyethyl)phosphine, 10 mM Mg(OAc)_2_, 50 mM potassium glutamate, 0.1 mM EDTA), and contained 1.5 nM DNA substrate substrate (see section on Forked Linear DNA substrates) 30 nM CMG, 30 nM MTC, 20 nM Pol ε, 10 nM RFC, 30 nM PCNA, 600 nM RPA, 5 mM ATP and 120 μM dNTPs, and where indicated 40 nM sgRNA and 20 nM dCas9-dL5 in a final volume of 20 μL. First, DNA was incubated with CMG and MTC for 2 min at 30°C followed by an additional 2 min with dCas9-dL5 and cgRNAs. Components except ATP and RPA were added and further incubated for 5 min at 30°C. Replication was initiated by addition of ATP and RPA. The reactions proceeded for the indicated amount of time at 30°C and were quenched with an equal volume of 2x stop solution (40 mM EDTA and 2% (*w/v*) SDS). DNA products were separated on a 1.3% (*w/v*) an alkaline agarose gel at 35 V for 16 h. Gels were backed with DE81 paper (GE Healthcare), dried by compression, exposed to a phosphorimager screen, and imaged with a Typhoon FLA 9500 PhosphorImager (GE Healthcare).

## *In vitro* single-molecule fluorescence microscopy

*In vitro* single-molecule microscopy was performed on an Eclipse Ti-E inverted microscope (Nikon, Japan) with a CFI Apo TIRF 100x oil-immersion TIRF objective (NA 1.49, Nikon, Japan), as previously described^6^. The temperature was maintained at 31°C (unless otherwise stated) by an electronically heated flow-cell chamber coupled to an objective heating jacket (Okolab, USA). NIS-elements was used to operate the microscope and the focus was locked through Perfect Focus System (Nikon, Japan). Images were captured using a Evolve 512 Delta EMCCD camera (Photometics, USA) with an effective pixel size of 0.16 µm. DNA molecules stained with 150 nM Sytox orange were imaged with a CW 568-nm Sapphire LP laser (200 mW max. output), and ET600/50 emission filter (Chroma, USA) at 0.76 W/cm^2^. dCas9-dL5–MGE complexes were imaged with a CW 647-nm OBIS laser (100 mW max. output), and 655LP emission filter (Chroma, USA) at 57.7 W/cm^2^.

## Preparation of flow-cells for *in vitro* imaging

Replication reactions were carried out in microfluidic flow-cells constructed from a PDMS flow chamber placed on top of a PEG-biotin-functionalized microscope coverslip as previously described^5, 6, 16, 17^. Once assembled, all surfaces of the flow-cell including connecting tubing were blocked against non-specific binding by introduction of 1 mL malic acid buffer (100 mM Na.maleate pH 7.5 and 250 mM NaCl) containing 1% (*w/v*) blocking reagent (Roche).

## Single-molecule rolling-circle blocking replication assays

The overall experimental scheme was to first form the dCas9-dL5–cgRNA1–DNA complex. Next, dCas9-dL5–cgRNA1–DNA complex was attached via the 5’-biotinylated flap-primed 2030-bp dsDNA circle bearing a 25-nt fork gap, to the surface via a biotin–streptavidin bond. Following a wash to remove unbound dCas9-dL5, replication was initiated by continuous flowing of reconstituted replisomes, ATP, dNTPs, and rNTPs and flow-stretching the DNA.

Specifically, 10 nM dCas9-dL5 was incubated with 200 nM cgRNA1 for ~5 min at 37°C in single-molecule imaging (SM) buffer (25 mM Tris-HCl pH 7.6, 10 mM Mg(OAc)_2_, 50 mM potassium glutamate, 0.1 mM EDTA, 10 mM dithiothreitol and 0.0025% (*v/v*) Tween20). The dCas9-dL5– cgRNA1 complex was then incubated with 100 pM replication template for a further 20 min at 37°C. The dCas9-dL5–cgRNA1–DNA complexes were adsorbed to the surface in SM buffer + 150 nM Sytox orange at 10 μL/min until an appropriate surface density was achieved. The flow-cell was then washed with 200 μL of SM buffer containing 50 mM NaCl. Following this replication was initiated — *E. coli* leading- and lagging-strand DNA replication reactions were carried out under the continuous presence of all proteins as previously described^6^. T7 leading- and lagging-strand DNA replication assays were carried out under the continuous presence of all proteins using previously described conditions^16^. All *in vitro* single-molecule rolling-circle blocking experiments were performed at least three times.

## Analysis of agarose gels of replication products

Agarose gel images were adjusted for brightness and contrast for clear visualisation using FIJI^18^. Blocked replication products were quantified in FIJI using in-house built plugins, by comparing the integrated intensity of bands between control and reaction lanes; the resulting percentages were then corrected for background and for the specified control.

## *In vitro* image analysis

Image analysis was performed in FIJI, using the Single Molecule Biophysics plugins (available at https://github.com/SingleMolecule/smb-plugins). Raw videos (.nd2 format) were converted into TIF files and flattened with the excitation beam profile as described previously^19^. For quantification of DNA product lengths, intensity projections were generated by summing 10 frames to reduce the contribution of transverse Brownian fluctuations of the DNA. Product length was determined by deconvolving the length of the rolling-circle substrate using the calibrated pixel size in bp (here, 1 pixel = 470 bp). Product length distributions were fit with a single-exponential decay (assuming a single rate-limiting step determining the end of an event). All distributions were made and fitted using MATLAB (Mathworks, USA).

## Constructs for live cell imaging

Plasmids: pdCas9dL5 was constructed by sub-cloning the *Not*I –*Xho*I insert from pJL001 into pJM1071. pJM1071 was a generous gift from the Woodgate laboratory^20^. Guide RNA plasmid pgRNA *recA*T 1 was constructed by inserting the *Nco*I –*Eco*RI insert into pBAD-myc-HisB. This insert carries the P104 promoter (Supplementary Table 1, bold)^21^, target sequence (Supplementary Table 1, underlined) and gRNA scaffold (supplementary Table 1, italicized).

Strains: The MG1655/pdCas9-dL5 pgRNA *recA* T1 strain was created by co-transformation with the two plasmids and selection using the appropriate antibiotics spectinomycin (50 μg/mL) and ampicillin (100 μg/mL) on plates. A single colony was grown out in LB media and a freezer stock was made by addition of 78 μL of DMSO to 1 mL of culture grown to mid-exponential phase at 37°C. MG1655 dCas9YPet was a generous gift of Johan Elf. The dCas9-Ypet/pgRNA *recA* T1 strain was created by transforming MG1655 dCas9-YPet with the pgRNA *recA* T1 plasmid and selected on LB plates containing 100 μg/mL ampicillin.

## Growth assays

Toxicity of MGE to cells was assayed in growth assays using a plate-reader. MG1655 cells carrying a pBAD-myc-HisB plasmid (amp^R^)^22^ were grown overnight at 30°C in EZ-glucose medium supplemented with 100 μg/mL ampicillin to obtain a stationary phase culture. Overnight culture (6 μL) was diluted into 6 mL of fresh EZ-glucose supplemented with ampicillin. Aliquots (5 x 1 mL) of the culture were then supplemented with either 0, 2, 20, 200, or 2000 nM MGE by diluting the dye 1000x from a corresponding series of MGE stocks. Aliquots (200 μL) of the culture were then plated in triplicate in a 96-well plate and cell growth was monitored by measuring absorbance at 600 nm of growing cell cultures every 20 min for 12 h. The average of the triplicate readings from one experiment are presented here. Experiments were repeated in EZ-glycerol medium and similarly no growth defects were observed (data not shown).

## Live cell imaging

Live cell imaging was performed on a custom built TIRF microscope^23^ equipped with a CW 647 nm (100 mW max output, OBIS, Coherent, CA, US set to at 57.7 W/cm^2^) and 514 nm (Sapphire, Coherent, CA, US set to 71 W/cm^2^) laser lines and a motorized stage. Cellular imaging was performed using highly inclined and laminated optical sheet imaging (HiLo) enabling illumination of bacterial cells immobilized on the surface using an inverted fluorescence microscope (Nikon Eclipse-Ti) equipped with a 1.49 NA 100x objective and a 512 x 512 pixel^2^ Andor iXon 897 EMCCD camera (Andor, US). NIS-Elements equipped with JOBS module was used to operate the microscope (Nikon, Japan).

The 514-nm laser light was directed through a 405/514/568 dichroic and ET535/30m emission filter (Chroma, Vermont, US); 647-nm laser light was directed through a 488/514/568/647 dichroic, and a 655LP emission filter (Chroma, USA). dCas9-YPet was imaged continuously with one hundred 100 ms exposures to 514 nm light. dCas9-dL5-MGE was imaged by exposing cells to one hundred 2 s exposures with CW 647-nm OBIS laser (100 mW max. output), and 655LP emission filter (Chroma, USA) at 57.7 W/cm^2^.

Photo-bleaching experiments were performed by exposing cells expressing an Mfd-dL5 (61 ROIs) or Mfd-YPet (254 cells) fusion to continuous laser irradiation while recording the emitted signal. To measure the photobleaching rate in cells, ROIs were drawn in cells and the total fluorescence intensity in each ROI was corrected for the background fluorescence. The fluorescence signal was then normalized by the maximum intensity (measured in the first frame). The normalized intensity curves as a function of time are presented in Supplementary Fig 1e.

## Flow cell construction

Custom flow-cells for live-cell imaging were constructed by gluing a (3-aminopropyl)triethoxy silane (Alfa Aesar, A10668, UK) treated coverslip (Marienfeld, Deckglaser, 24 x 50 mm No. 1.5, Germany) to a quartz piece (Proscitech, Australia) using double-sided sticky tape (970XL ½ X 36yd, 3M, United States) to create a channel, and sealed with epoxy. Quartz pieces were designed to provide an inlet and outlet tubing (PE-60, Instech Labs).

## Preparation of cells prior to imaging

Prior to imaging, cells were revived from a –80°C freezer stock by resuspension in LB liquid media overnight (500 μL) and shaking at 1000 rpm at 30°C on an Eppendorf thermomixer C (Eppendorf, Australia). On the following day, cells were diluted in EZ rich defined medium supplemented with 0.2% *w*/*v* glucose as the carbon source (EZ-glucose, Teknova, CA, US). Cells were inoculated 1:1000 in 500 μL of EZ-glucose and shaken for 4 h or until the *A*_600_ was ~0.2. Cells were then introduced into the flow cell and allowed to settle on to the surface of the coverslip. Finally, the cell suspension was switched with fresh, aerated growth medium supplemented with 20 nM MGE before starting the experiment. Imaging was performed at 30°C on a heated stage coupled to a heated objective (OKO Labs).

## Image analysis

The beam profile was obtained by creating an average projection of fluorescence images collected in each imaging session for each channel followed by Gaussian filtering with a radius of 50 pixels. Images were flattened using the beam profile and a dark count of 375 for the 514 nm channel and 1200 for the 647 nm channel. Image processing used custom software in FIJI^24^.

## Supplementary Information

**Supplementary Fig. 1:**
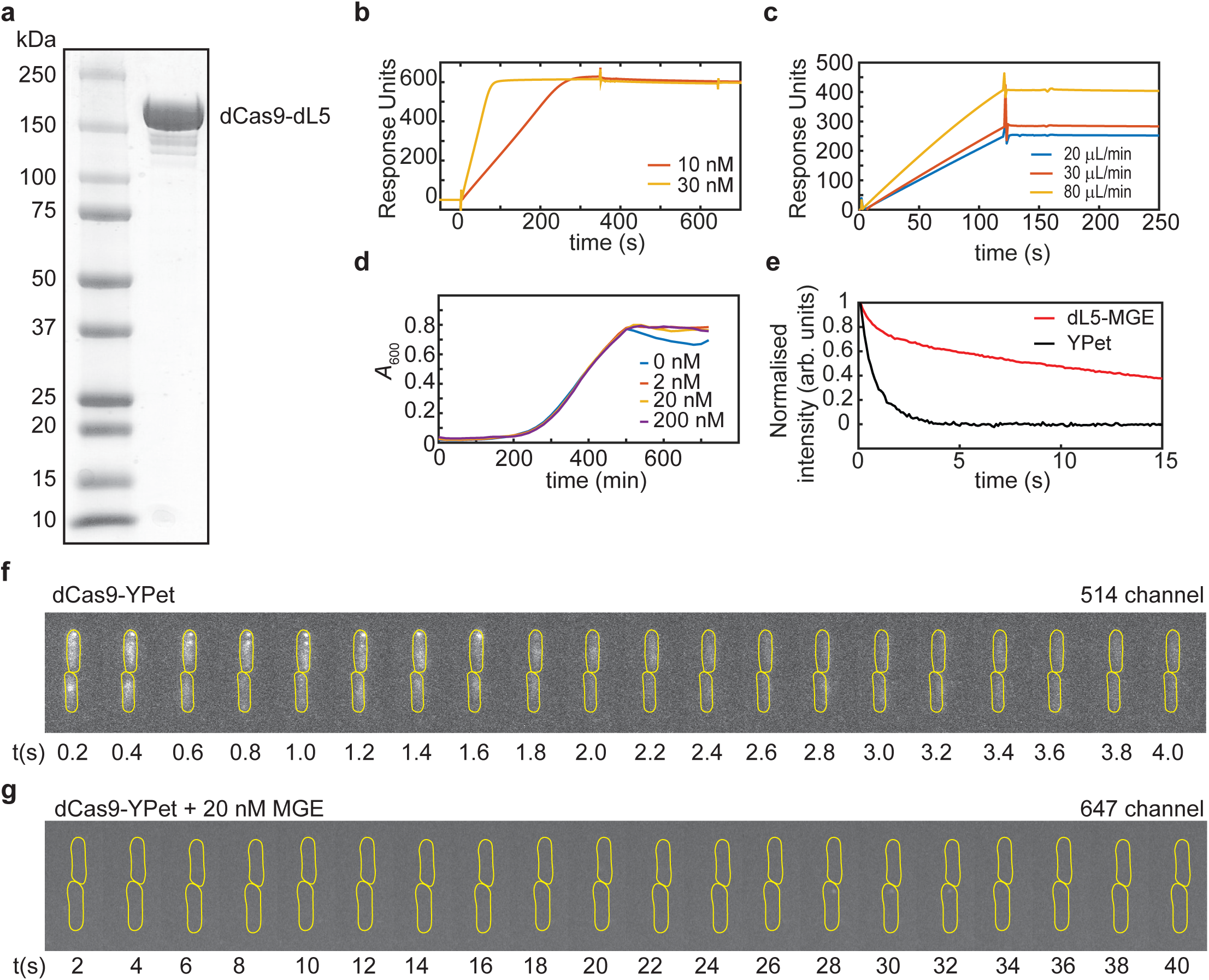
Characterization of dCas9-dL5. **a.** Coomassie stained 4–20% SDS-PAGE of purified dCas9-dL5. **b.** Sensorgrams showing binding to and dissociation of 10 and 30 nM dCas9-dL5-cgRNA from 83- mer dsDNA substrate immobilized on an SPR chip. c. Sensorgrams monitoring the association of dCas9-dL5-cgRNA (10 nM) injected at three different flow rates (20, 30 and 80 µL/min) onto 83-mer dsDNA containing target sequence immobilized on an SPR chip. The linearity and difference in responses indicates mass transfer limitation. **d.** Growth curves of MG1655/pBAD-myc-HisB at 30°C in EZ-glucose growth medium supplemented with the indicated amount of MGE dye. **e.** Normalized intensity traces (average and standard error of mean) from YPet (blue curve, 254 cells) and dL5-MGE (red curve, 61 ROIs) from Mfd-Ypet and Mfd-dL5 fusions in MG1655. **f.** Time-series acquisition of dCas9-YPet targeted to the *recA* locus in MG1655. Cells were excited with 514 nm laser light and emission was collected in the 535/30 nm channel. **g.** Time-series acquisition of dCas9-YPet targeted to the *recA* locus in MG1655 in EZ-glucose medium supplemented with 20 nM MGE. Cells were excited with 647 nm laser light, and the emission was collected using a 650 LP filter. MGE alone does not exhibit foci in the absence of the dL5 tag.

**Supplementary Fig. 2:**
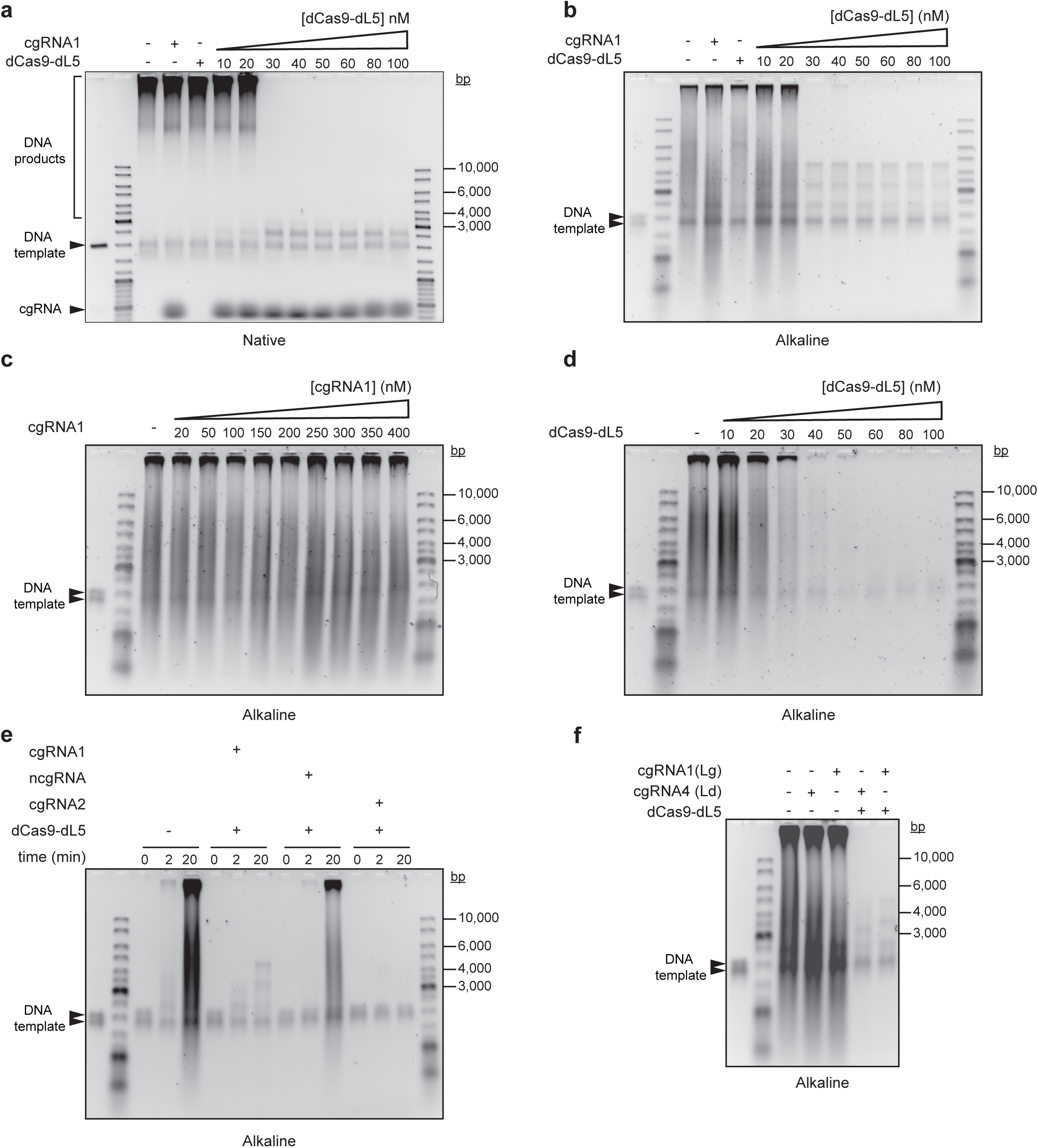
Target-bound dCas9-dL5 site-specifically arrests *E. coli* DNA synthesis. **a.** Target-bound dCas9-dL5 arrests *E. coli* leading strand DNA synthesis. Unless otherwise specified reactions contained 400 nM cgRNA1 and 20 nM dCas9-dL5. At concentrations below 20 nM, dCas9-dL5 does not completely arrest leading-strand DNA synthesis. **b.** Target-bound dCas9-dL5 arrests *E. coli* leading- and lagging-strand DNA synthesis. Unless otherwise specified, reactions contained 400 nM cgRNA1. At concentrations below 20 nM, dCas9-dL5 does not completely arrest leading- and lagging-strand DNA synthesis. **c.** High concentrations of complementary gRNAs do not inhibit *E. coli* leading- and lagging-strand DNA synthesis. Unless otherwise specified, 50 nM dCas9-dL5 was used for all reactions. **d.** dCas9-dL5 alone does not site-specifically inhibit *E. coli* leading- and lagging-strand DNA synthesis. Non-specific inhibition is observed at high concentrations of dCas9-dL5 alone (see Supplementary note 5, Fig. 2a and summary in Fig. 2h). **e.** Only dCas9-dL5 programmed with complementary gRNAs specifically arrests *E. coli* leading- and lagging-strand DNA synthesis. Reactions contained 50 nM dCas9-dL5 and 400 nM gRNAs. Reactions were initiated at 30°C and aliquots were removed and quenched at 0, 2, and 20 min time points. **f.** *E. coli* leading- and lagging-strand DNA synthesis arrest by target-bound dCas9-dL5 is not strand specific. Unless otherwise specified reactions contained 400 nM cgRNAs and 50 nM dCas9-dL5. Lg denotes cgRNA targeted to the lagging strand, and Ld denotes cgRNA targeted to the leading strand. All panels show photographic negative images of gels that had been stained with SYBR-gold nucleic acid stain.

